# Identification of RNA-associated peptides, iRAP, defines precise sites of protein-RNA interaction

**DOI:** 10.1101/456111

**Authors:** Lauri Peil, Sakharam Waghmare, Lutz Fischer, Michaela Spitzer, David Tollervey, Juri Rappsilber

## Abstract

The identification of ever-increasing numbers of RNA species has underlined the importance of robust characterization of bona fide sites of protein-RNA interaction. UV crosslinking can be used to identify precise RNA targets for individual proteins, transcriptome-wide. Here we sought to generate reciprocal data, identifying precise sites of RNA-binding proteome-wide. The resulting technique, identification of RNA-associated peptides (iRAP), was used to locate 1331 unique RNA-interaction sites at single amino acid residue resolution in 324 *S. cerevisiae* proteins. Our identified RNA-interaction sites in characterized RNA-protein complex agree well with available high-resolution structures. In known RNA-interacting protiens, 317 sites fall into known and suspected RNA-interaction domains while only 21 sites fall into other annotated sequence features. Strikingly, 993 of the sites identified fall into protein regions that lack any recognizable protein domain structure or annotated sequence features. This suggests that, despite binding RNA *in vivo*, many of these proteins will not have defined functions in RNA metabolism.

**Highlights:**

Method identifies precise RNA-binding sites across proteome
1331 sites identified that contact RNA in 324 proteins in *S. cerevisiae*
RNA binding takes place predominantly outside known protein domains

## Introduction

Interactions between RNA and proteins play key roles in many aspect of cell metabolism. However, the identification of protein-RNA interaction sites has long been challenging, particularly in living cells. Individual protein-RNA interactions can be characterized, if known, by mutagenic and biochemical approaches but this has always been labor intensive. The difficulty is compounded by the fact that many interactions do not fall within characterized interaction domains, and even apparently well-characterized RNA-binding domains can show multiple modes of RNA interaction (Clery, Sinha et al., 2013), making detailed predictions less reliable.

For large-scale characterization of protein-RNA interactions, a significant advance was the development of RNA immunoprecipitation (RIP) with or without formaldehyde crosslinking, allowing the identification of RNAs associated with target proteins, although not the site of association (Gilbert, Kristjuhan et al., 2004, Gilbert & Svejstrup, 2006, Huang, Johansson et al., 2005, Hurt, Luo et al., 2004, Motamedi, Verdel et al., 2004, Niranjanakumari, Lasda et al., 2002). Subsequently, UV crosslinking approaches were developed that allow accurate identification of the binding sites for individual proteins on RNA molecules, transcriptome-wide (Bley, Qi et al., 2011, Doneanu, Gafken et al., 2004, Granneman, Kudla et al., 2009, Granneman, Petfalski et al., 2010, Maly, Rinke et al., 1980, Mital, Albrecht et al., 1993, Rhode, Hartmuth et al., 2003, Urlaub, Hartmuth et al., 2000, Van Nostrand, Pratt et al., 2016, Wagenmakers, Reinders et al., 1980).

The reciprocal analyses of proteins that are bound to RNA have been more difficult to develop, at least in part because proteomic approaches do not provide the amplification offered by PCR. However, recent reports have successfully used UV crosslinking and RNA enrichment to identify many or all poly(A)^+^ RNA binding proteins present in human cells and other systems (Baltz, Munschauer et al., 2012, Castello, Fischer et al., 2012, Kwon, Yi et al., 2013). This technique was an important advance but, like RIP, identifies the species involved but not the site of interaction, and is limited to mature mRNAs. To identify the total RNA-bound proteome, the approach of 5-ethynyluridine (EU) labeling of RNAs followed by biotin ligation using the click reaction (RICK) was recently developed (Bao, Guo et al., 2018, Huang, Han et al., 2018). In addition, MS analyses have been developed to identify the precise amino-acid at the site of RNA-protein crosslinking (Kramer, Sachsenberg et al., 2014).

Here, we sought to develop a ready means for the identification of the peptide, and indeed the amino acid that is crosslinked to RNA during UV irradiation in living cells. This is based on the identification of the specific mass difference resulting from the single residual nucleotide that remains associated with the peptide following complete hydrolysis of the RNA and partial digestion of the protein. Here we report the application of this technique to precisely map the RNA-binding sites of hundreds of yeast proteins following crosslinking in actively growing cells, validated by using available high-resolution structures for yeast ribosome and individual protein-RNA complexes, together with genome-wide analysis of RNA targets of the novel RNA-binding protein enolase. This approach should be widely applicable for the reliable and accurate characterization of the protein-RNA interactome in many systems.

## Results

### Development of iRAP protocol

To identify proteins that directly bind to RNA and their attachment sites to RNA, we first covalently attached RNA to its associating proteins by UV crosslinking in actively growing cells. Our initial rationale was that RNA purification under stringent washing conditions should retain only covalently attached proteins or peptides and thus reveal the identity of those proteins that directly contact to RNA. When proteins are then digested by a specific protease such as LysC or trypsin, only peptides attached to the RNA and thereby harboring the actual linkage sites, can be purified together with RNA. Finally, RNA degradation should leave a single nucleotide attached to peptides and these peptides could then be identified by mass spectrometry, a process that would also reveal the exact site of nucleotide-modification. In this way, protein identity and RNA-attachment point would be revealed simultaneously for all proteins.

In establishing the purification protocol for RNA-associated peptides, various conditions were tested, with the best results being obtained using a CsTFA density gradient. This separation is based on the previous observations that RNA is denser than proteins, with RNA-protein complexes showing an intermediate density.

To assess whether the purification of RNA-protein complexes recovered only covalently bound conjugates, a SILAC experiment was performed to quantify protein recovery in the presence and absence of UV crosslinking. Yeast strains that were auxotrophic for lysine (*lys2*Δ*0*) were grown in the presence of lysine or [^13^C_6_]-lysine. We mixed unlabeled, non-crosslinked lysate with isotope labeled *in vivo* crosslinked lysate prior to fractionation. Unexpectedly, even after stringent RNA purification (lysis in the presence of urea and/or guanidinium thiocyanate, RNA purification through CsTFA gradient), the majority of identified non-modified peptides showed no enrichment in the cross-linked sample (Figure S1A, S1B). This suggests that the identification of RNA-bound proteins based solely on co-precipitation with RNA even by help of crosslink stabilization and stringent washing may not be a fully reliable indicator of *in vivo* interactions. We therefore focused on identifying the subset of peptides of these proteins that had been covalently linked to nucleic acid, since these links can arise only *in vivo* during the irradiation step. Indeed, no RNA-modified peptides were identified in the absence of cross-linking (see below).

To identify RNA-peptide conjugates, we used the final workflow outlined in Fig 1A. Briefly, 2 l cultures of actively growing yeast were UV-irradiated at 254 nm as previously described (Granneman et al., 2009). Recovered cells were lysed and proteins digested under denaturing conditions. RNA was purified from the digestion mixture, taking along any covalently attached peptides, purified RNA was hydrolyzed, RNA-modified peptides were separated from nucleotides, desalted and analyzed via LC-MS/MS. We named this approach identification of RNA associated peptides (iRAP). Detailed variations of the recommended iRAP workflow are given in Supplemental Data.

**Figure 1.**
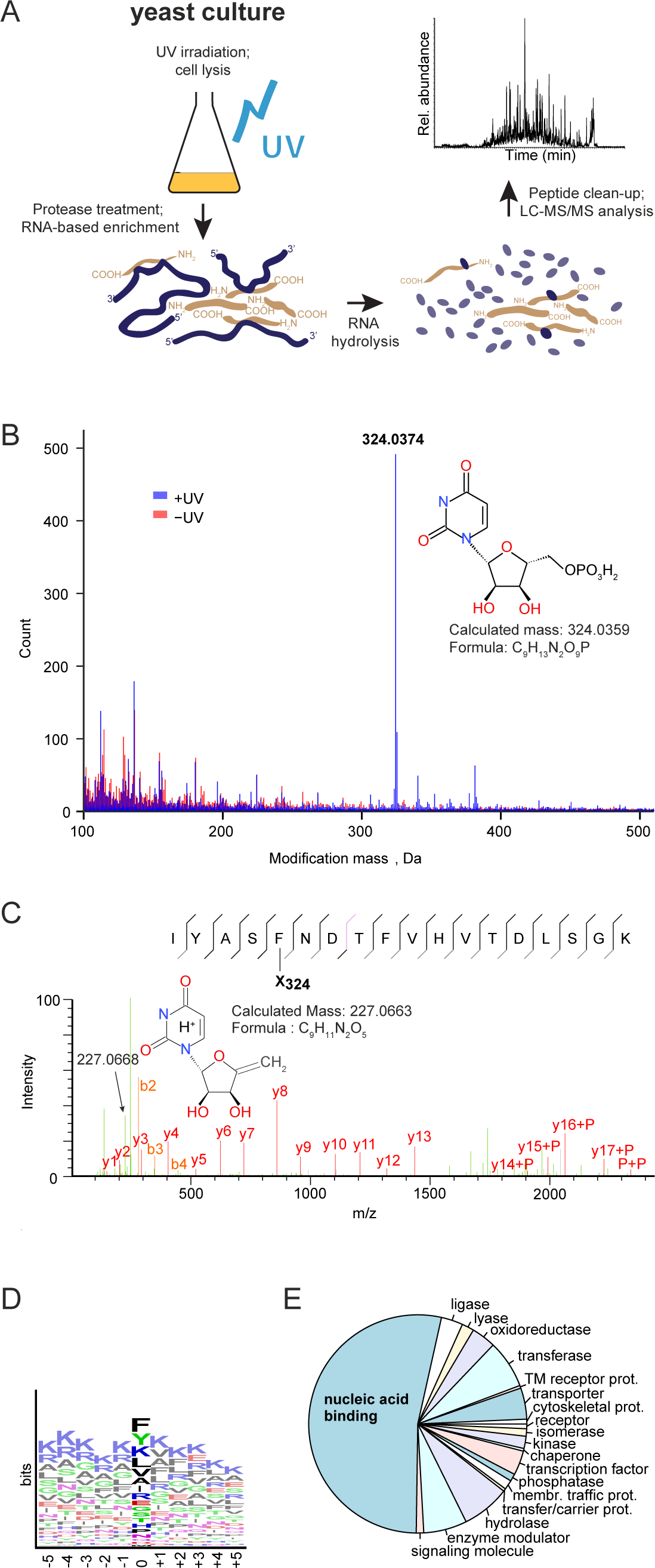
UV irradiation induces protein:RNA crosslinks via uridine. A. Workflow used for identifying peptides crosslinked to RNA. Yeast culture was UV-irradiated, cells lysed under denaturing conditions and RNA-crosslinked proteins digested with LysC. Total RNA (together with RNA-bound peptides) was extracted, RNA was hydrolyzed, peptides were cleaned up and analyzed by LC-MS/MS. B. UV-irradiation forms a covalent peptide modification that corresponds to uridine. Peptide modification masses from xiOMS open modification search are plotted for UV-irradiated (blue) and non-irradiated (red) samples. UV-specific modification with a mass of 324.0374 Da corresponding to uridine monophosphate is shown. C. Annotated fragmentation spectrum of an UMP-modified peptide. Amino acid sequence and modification site is shown for an UMP-modified peptide, together with annotated b and y fragment ions on the spectrum. Also indicated is a potential reporter ion with a m/z of 227.0668, together with its proposed structure. D. Sequence logo representation of UMP-modified amino acids in identified peptides. All identified peptide:RNA linkage sites with a 324.0374 Da modification in an initial data analysis were analyzed for amino acid enrichment using WebLogo 2 tool at weblogo.berkeley.edu. E. PANTHER Classification System analysis of RNA-linked proteins according to their ProteinClass annotations.

### Identification of RNA-binding sites in proteins

An initial dataset, generated from poly(A)^+^-enriched RNA-protein complexes, was searched using in-house developed Xi database search platform (ERI, Edinburgh) with an open modification search option (xiOMS), to identify any UV-specific modifications. xiOMS identifies peptides in a modification-independent manner, allowing a search for unknown and unexpected modification masses. This search revealed an abundant unknown UV-specific modification with a mass of 324.0374 Da (Fig 1B). None of the peptides carrying this modification were found by SILAC in the non-UV-irradiated control. This indicates strongly that 324 Da is a UV-dependent modification of peptides. Elemental composition simulation proposed this modification to be C9H13N2O9P (324.0359 Da, 4.7 ppm), which corresponds to uridine monophosphate (UMP). Interestingly, this was the only nucleotide-derived modification we were able to identify, we observed no evidence for peptide cross-links with A or G or C. We also observed a modification of 244.0713 Da, consistent with the mass of uridine (244.0695 Da, 7.2 ppm), but this was not used for further analysis due to its low occurrence.

Fig 1C shows an example of an UMP-modified peptide identified in xiOMS, with the exact modification site determined by full b- and y- ion ladders. In further support of the identification of the modification as UMP, we also observed the presence of a potential reporter ion at m/z 227.0668, a mass that fits with the elemental composition of uridine after elimination of H3PO4 during fragmentation (227.0662 Da, −2.6 ppm) (Fig 1C). Next, we generated sequence logos with the modified peptides to determine whether some UMP-modified amino acids are overrepresented (Fig 1D). There was an over-representation of aromatic amino acids phenylalanine and tyrosine, together with lysine, leucine, valine, alanine and isoleucine. This is in agreement with Phe and Tyr frequently being found as RNA-interacting residues in RRMs (Clery, Blatter et al., 2008, Hobor, Pergoli et al., 2011, Teplova, Song et al., 2010, Tsuda, Kuwasako et al., 2009). As a result, we used these seven amino acids together with previously described tryptophan (Bley et al., 2011, Katouzian-Safadi, Laine et al., 1991, Reeve & Hopkins, 1980) to define new modifications in the MaxQuant search engine (Cox & Mann, 2008). All of the acquired data was subjected to large-scale data analysis using two different search engine platforms; either MaxQuant or using an in-house developed, Xi database search platform (ERI, Edinburgh) with a target modification search option (xiTMS) (see Methods and Supplemental Information). The main difference between these two search platforms was that in MaxQuant UMP modification was defined to be specific to the most frequently identified amino acids, whereas xiTMS was targeted to UMP modifications with no amino acid specificity, thereby making these two approaches complementary to each other.

Combining all results from xiOMS, xiTMS and MaxQuant searches, we identified 324 unique RNA-linked proteins in total, with 1331 unique linkage sites (Table S1 and Table S2, respectively). Ribosomal proteins, including mitochondrial ribosomal proteins, comprised about 20% of RNA-linked proteins and 40% of unique linkage sites, with 68 proteins and 526 linkage sites identified (Table S1 and Table S2). For 28 ribosomal proteins it was not possible to distinguish between the A and B isoforms, since these are identical for the peptides found to be crosslinked to RNA.

Using PANTHER (Protein ANalysis THrough Evolutionary Relationships), a classification system designed to classify proteins (and their genes), (Mi, Muruganujan et al., 2013a, Mi, Muruganujan et al., 2013b), 281 out of 324 identified RNA-crosslinked proteins could be annotated based on their Protein Class annotation (Fig 1E, Table S1). 149 proteins (53%) were classified as ‘nucleic acid binding’, with additional 40 proteins classified either as “transferase” or as “hydrolase”.

### Ribosome as a proof of principle for iRAP

Ribosomal proteins in which RNA crosslinks were mapped are shown on the yeast 80S ribosome structure (Fig 2A). Of the 73 ribosomal proteins in the yeast 80S ribosome structure (PDB 3U5B|3U5C|3U5D|3U5E) (Ben-Shem, Garreau de Loubresse et al., 2011), 56 were identified as UMP-modified, with a total of 436 linkage sites (Table S3). As an example, Thr20 of Rpl2A and its neighboring uridine are indicated in a zoom-in (Fig 2A), demonstrating the close proximity of the amino acid and the nucleotide. To test whether this is also true for other ribosomal linkage sites, we analyzed the distributions of shortest distances from each amino acid to the closest uridine in the yeast 80S ribosome structure (Fig 2D). Fig 2B compares the shortest distance between each amino acid and its closest uridine (indicated in magenta) with the shortest distance between each RNA-linked amino acid and its closest uridine. RNA-linked amino acids in the yeast ribosome were generally closer to uridine residues than all amino acids, with the two populations showing highly significantly differences (P value 1.5×10^-62^) as determined by a Kruskal-Wallis test (Kruskal & Wallis, 1952). Note, however, that complete coincidence between the observed crosslinks and the crystal structure is not expected, since the crosslinking was performed on active, functional ribosomes, whereas inactive ribosomes were purified from glucose-starved cells for crystallization (Ben-Shem et al., 2011).

**Figure 2.**
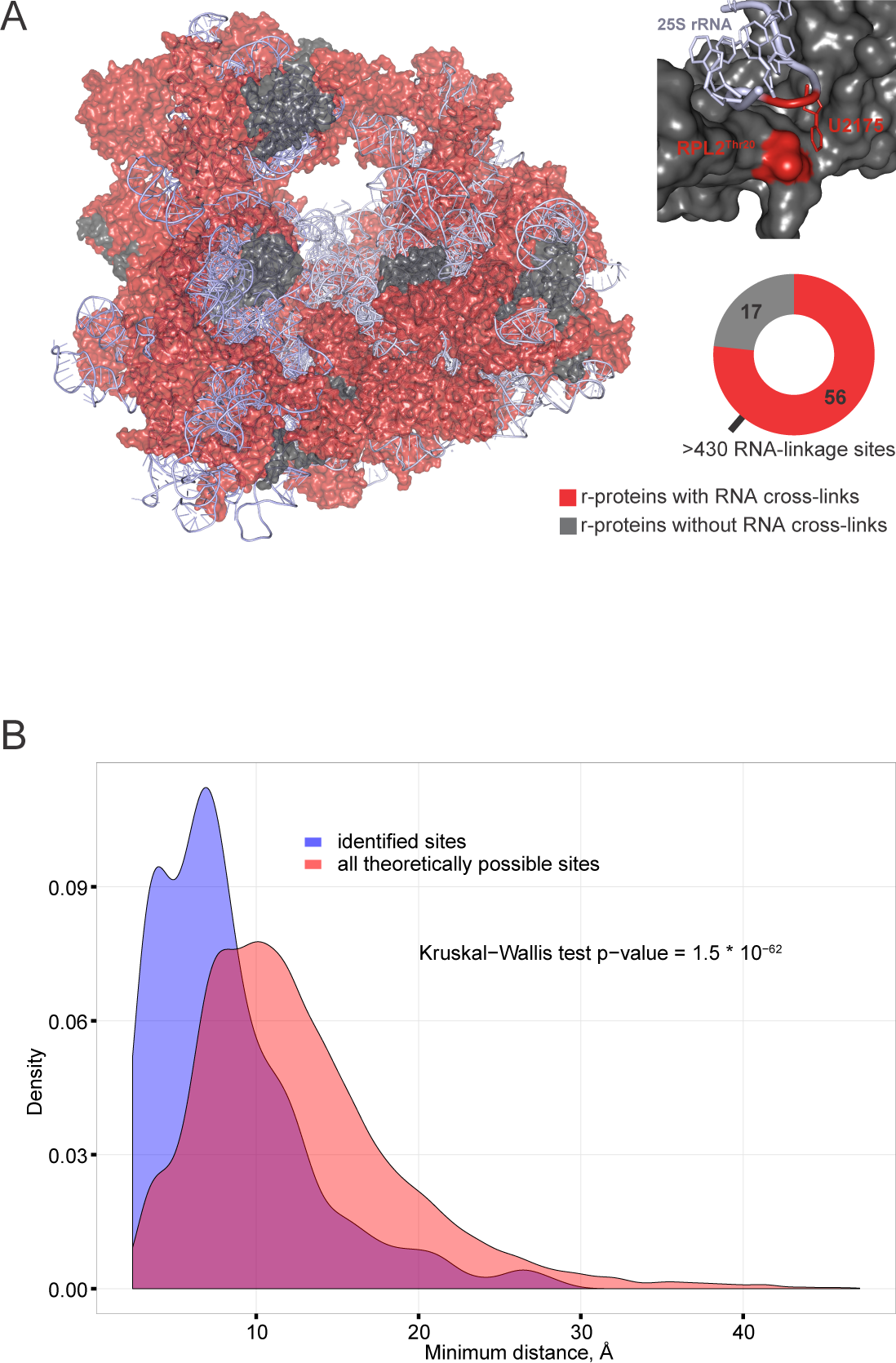
Yeast ribosome structure supports the protein:RNA crosslinks identified. A. Yeast 80S ribosome crystal structure (PDB 3U5B|3U5C|3U5D|3U5E). Ribosomal proteins with evidence for RNA crosslinks are shown in red, non-crosslinked proteins are shown in dark gray and ribosomal RNA in light gray. In total, 56 RNA-linked ribosomal proteins with more than 500 RNA linkage sites were identified. In a zoom-in box ribosomal protein RPL2 and part of 25S rRNA are shown in gray, with Thr20 residue of RPL2 and its closest uridine highlighted in red. C. Distances between identified RNA-linked amino acids to their closest uridine in the yeast 80S ribosome are shorter than all theoretical distances. For distance measurements, only amino acid side-chain atoms and uracil atoms were considered, and only the shortest distance was recorded for each amino acid:uracil pair. The distribution of distances for identified RNA linkage sites is shown in blue, the distribution of distances for all chemically possible linkage sites is shown in red. The significance of the difference between the two distributions was determined by Kruskal-Wallis test.

### Mapping of protein-RNA cross-linking sites to protein features

Next, we analyzed all identified RNA-crosslinked proteins with regard to their UniProtKB annotated sequence features. Only sequence features for which at least two independent RNA crosslinks were identified were included (Fig. 3A). By far the most abundant sequence feature was the RNA-recognition motif (RRM), with RNA crosslinks localized to 37 individual RRM domains in 27 proteins. Other sequence features were represented with lower numbers, with only Pumilio having RNA linkage sites falling into seven individual features in three proteins. As expected, almost all of the annotated sequence features are involved in RNA binding or metabolism, with an exception of the “Protein kinase” motif, which was recovered two times.

**Figure 3.**
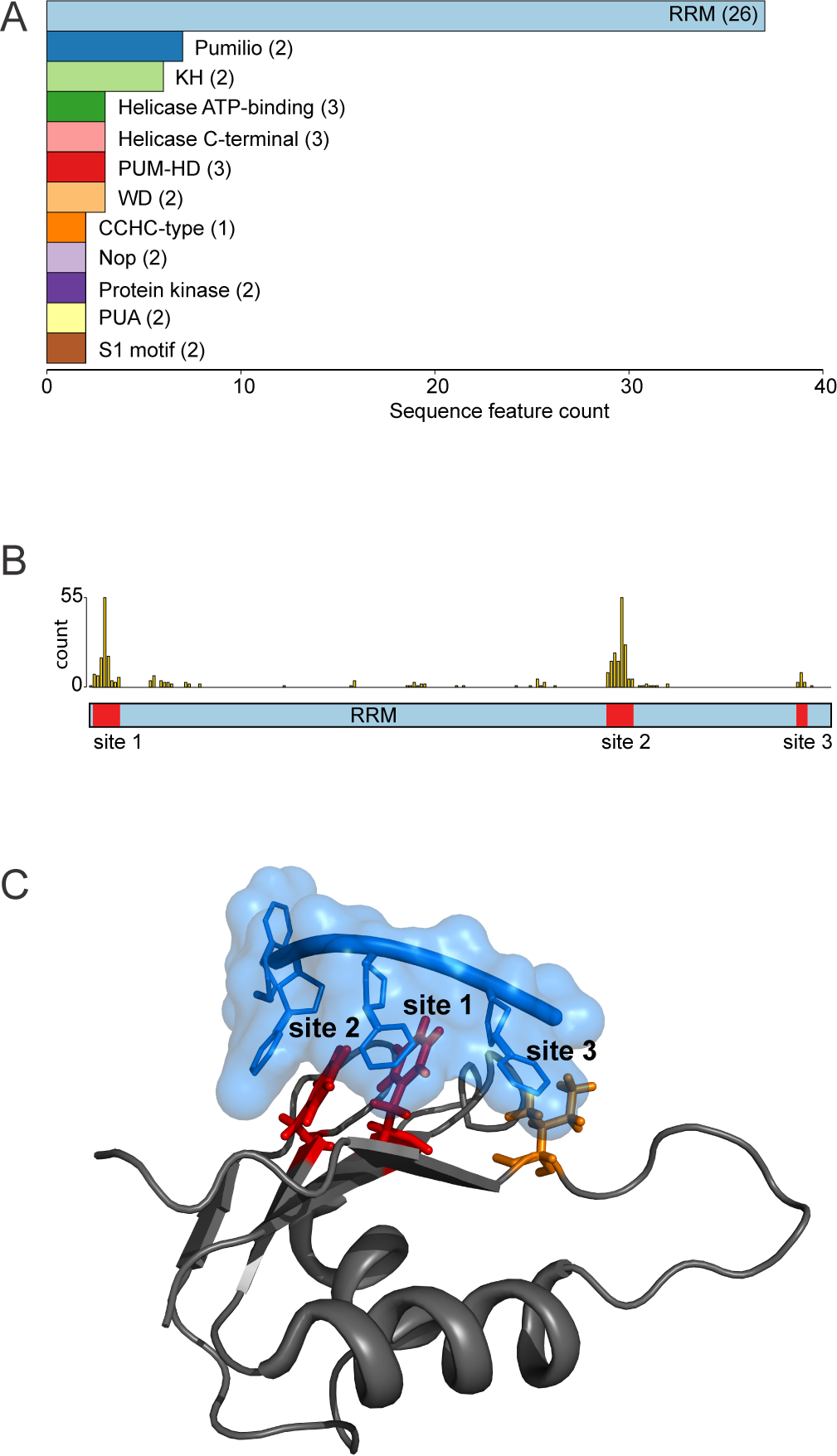
RNA linkage sites identified in proteins are localized to annotated RNA-interacting domains. A. Protein sequence feature analysis. All identified 324 RNA-linked proteins were analyzed for their UniProtKB-annotated protein sequence feature content. Only sequence features with evidence for two or more RNA linkage sites are shown, number of unique proteins for each sequence feature is shown in brackets. B. Histogram of identified RNA linkage sites in RRM domains. All RRM domains with identified RNA linkage sites were aligned and linkage sites were counted and plotted on a histogram. Three main linkage site regions are indicated. C. Crystal structure of Nab3 RRM domain in complex with UCUU (PDB 2L41). Nab3 RRM domain is displayed in gray, UCUU in blue and identified RNA-linked amino acids in red. In addition, a third potential linkage site is shown in orange, based on RRM domain sequence homology; all three linkage sites are numbered as in panel B.

Since the RRM domain was the best represented of all annotated sequence features and well documented in regard its RNA binding properties (Clery et al., 2008), we assessed the distribution of RNA linkage sites within RRM domains. As shown in Fig 3B, identified RNA linkage sites fall into three distinct regions of the RRM domains – N-terminus, central region and C-terminus. This finding is in very good correlation with an in-solution NMR structure (PDB 2L41) (Hobor et al., 2011) of an RRM domain from Nab3 in complex with an oligonucleotide UCUU reflecting its consensus binding site (Fig 3C). Two of the linkage sites (displayed in red) were identified in Nab3 in this study. We identified the third linkage site (displayed in orange) in other RRMs, and positioned it based on sequence alignment.

The Pumilio RNA-binding domain was also well represented among the RNA-binding peptides, and crosslinked residues were in good agreement with predicted contact sites. As an example, Figure 4 shows the crystal structure of the Pumilio domain of Puf3; identified RNA-linked amino acids are in close proximity to RNA nucleotides. Other known RNA binding domains were also well-represented in our data, but the lack of suitable high-resolution structures did not allow us to visualize RNA-linkage sites for confirmation.

Our set of identified RNA-linked proteins contained 33 unique UniProtKB-annotated sequence features, of which 21 were assigned as RNA-binding (Table S4). In total, 326 of 767 non-ribosomal protein linkage sites were mapped to 81 separate UniProtKB-annotated sequence features of 63 proteins (Table S2). Of those 326 linkage sites, 306 fall into known RNA binding domains in 53 different proteins. Another 204 non-ribosomal protein RNA linkage sites were assigned to 85 proteins that had UniProtKB-annotated sequence features but none of these linkage sites were mapped to any of those sequence features. Finally, 257 RNA linkage sites were assigned to 110 proteins that lacked any UniProtKB-annotated sequence features. We also analyzed low complexity regions in the identified proteins using the SEG program (Wootton & Federhen, 1996). 81 linkage sites fall into low complexity regions of which 69 are linkage sites that we could not map to annotated protein features.

We conclude that crosslinks identified in known RNA-binding proteins are generally located within the expected RNA-binding domains. However, crosslinks in proteins not predicted to bind RNA are generally in regions with no annotated structural domains.

## Discussion

Most previous UV-crosslinking studies were carried out with DNA, rather than RNA (for reviews see (Meisenheimer & Koch, 1997, Steen & Jensen, 2002, Williams & Konigsberg, 1991)). All native nucleobases have absorption range of 250-270 nm, meaning that any nucleic acid can be excited and will react with proteins (Meisenheimer & Koch, 1997, Williams & Konigsberg, 1991). Reported yields for *in vitro* protein:nucleic acid crosslinks are around 2-15%, (Steen & Jensen, 2002, Williams & Konigsberg, 1991), and in yeast cells we saw RNA crosslinking yields of up to around 5%. These levels suggested that it should be possible to recover sufficient material to identify exact amino acids within peptides that are UV crosslinked to RNA. However, crosslinking reaction mechanisms are not fully understood, making the products hard to predict (Shetlar, 1980, Steen & Jensen, 2002). Despite this, we were able to identify UV-specific peptide modification with a mass of 324 Da, corresponding to the addition of a uridine monophosphate, without any losses of functional groups or atoms. It has been noted previously that in the photochemical reactions involving thymidine and tyrosine, the masses of the adducts formed were found to be equal to the sum of the masses of individual species (Connor, Falick et al., 1998).

It was unexpected that only crosslinks with uridine were found, since CRAC analysis had identified all four other nucleotides as RNA interaction sites (Granneman et al., 2010). The basis of this specificity in the MS analysis is unclear, but we note that a similar phenomenon was previously reported an MS analysis of nucleotides recovered bound to peptides (Kramer et al., 2014). *In vitro* studies determined that the UV crosslinking efficiency of nucleotides can be ranked as dT >> dC > rU > rC, dA, dG (Hockensmith, Kubasek et al., 1986, Paradiso & Konigsberg, 1982, Williams & Konigsberg, 1991), consistent with our findings. In addition, UV-induced protein:nucleic acid complexes may differ in their stability to acids, bases, or denaturants (Shetlar, 1980). UV-induced protein:nucleic acid complexes are stable to high concentrations of salts and denaturants (Williams & Konigsberg, 1991). In contrast, incubation in acetic acid/methanol solution, similar to Coomassie fixing solution, disrupted up to 75% of a covalent gp32:DNA complex (Hockensmith, Kubasek et al., 1991). It therefore seems likely that other nucleotides within RNA do crosslink with proteins but are not abundant or stable enough to be detected under the conditions used by us during analysis. This may limit the recovery of proteins that specifically interact with RNA binding sites that lack uracil. However, it does not appear to be major problem for reciprocal CLIP-related assays.

Most amino acids can form photo-adducts with oligo-DNA or RNA sequences when single amino acids are added to the reaction, with Cys, Lys, Phe, Trp, Tyr, His, Glu and Asp being the most reactive amino acids towards DNA (Shetlar, 1980, Shetlar, Carbone et al., 1984, Shetlar, Christensen et al., 1984, Shetlar, Home et al., 1984). Based on previous analyses (Bley et al., 2011, Connor et al., 1998, Doneanu et al., 2004, Maly et al., 1980, Williams & Konigsberg, 1991, Zwieb & Brimacombe, 1979), aromatic amino acids would also be expected to be among the most reactive with intact proteins, and this was found in our analyses. In previous analyses, positively charged (Lys, Arg) or polar amino acids (Asn, Cys, Gln, His, Ser, Thr, Tyr, Trp) were largely implicated in interactions with nucleic acids, via either nitrogen bases or phosphate group (Ellis, Broom et al., 2007, Lejeune, Delsaux et al., 2005, Treger & Westhof, 2001) whereas we saw enrichment for hydrophobic amino acids (Ile, Leu, Val, Ala). As above, the restriction of the MASCOT database search to 7 amino acids may have led to the exclusion of a number of crosslinked proteins from the final list, which is unlikely to be exhaustive.

Structures of protein-RNA complexes deposited in the Protein Data Bank have been used previously to analyze interactions between amino acids and nucleotides (Ellis et al., 2007, Lejeune et al., 2005, Treger & Westhof, 2001). Interestingly, in all three studies it was found that Trp and Phe, together with Ala, Val, Ile and Leu were disfavored for interacting with RNA whereas these six amino acids were among the over-represented amino acids in our study. One can argue that available structures represent only a subset of all interactions and conclusions are therefore biased. An alternative explanation could be that whilst hydrophobic amino acids are not directly involved in RNA binding in studied structures, they are still in close enough proximity to be UV-cross-linked, especially considering that we found evidence for only crosslinks with U. Another potentially interesting finding was that RNA and DNA behave differently regarding protein interactions – most protein-DNA interactions involve phosphate atoms while protein-RNA involve more frequently base edge and ribose atoms (Lejeune et al., 2005). This knowledge may prove to be useful for future studies on protein-DNA interaction sites.

The majority of ribosomal proteins were identified with RNA crosslinks, including several mitochondrial r-proteins. Since the crystal structure of the yeast ribosome is available (Ben-Shem et al., 2011), distances could be calculated from each RNA-linked amino acid to the nearest uracil residue. Consistent with “zero-length” crosslinking being induced by UV, the RNA binding sites were generally located in close proximity to uracil residues. Some discrepancies were observed, but the ribosomes used for crystallization were extracted from glucose-starved, non-growing cells, and may differ in some details from the ribosomes in the actively growing cells used here. Moreover, since we did not identify which RNA species was crosslinked to each protein, some RNA interactions identified in our ribosomal proteins may be formed with the tRNA, mRNA or other RNA species, rather than the rRNA, thereby giving longer distances than predicted from the crystal structure data.

As expected, the non-ribosomal proteins that crosslinked to RNA were strongly enriched for known RNA-binding motifs. The most abundant was the well-characterized RRM domain, and the binding sites identified in these domains were in good agreement with the crystal structures of RRM-RNA complexes (Clery et al., 2008, Hobor et al., 2011, Teplova et al., 2010, Tsuda et al., 2009). Several other known nucleic acid binding motifs were also recovered, including Pumilio, KH and WD domains (Neer, Schmidt et al., 1994, Smith, Gaitatzes et al., 1999, Spassov & Jurecic, 2003, Valverde, Edwards et al., 2008), and helicase-specific domains (Tanner & Linder, 2001). As for RRM domains, RNA-linkage sites identified in Pumilio domains were in good agreement with the crystal structure of a Pumilio-RNA complex (Zhu, Stumpf et al., 2009).

Several recent analyses have used mass spectrometry to identify proteins that are recovered in association with RNA following UV crosslinking in human and other cells (Baltz et al., 2012, Bao et al., 2018, Castello et al., 2012, Huang et al., 2018, Kramer et al., 2014, Kwon et al., 2013). A striking finding from these reports and our work is that large numbers of proteins identified lack previously known RNA-binding activity or recognizable RNA-binding motifs. The recovery of multiple metabolic enzymes led to the hypothesis that cross talk between RNA and intermediary metabolism might be prevalent (White, 1976). Consistent with these results, we identified several metabolic enzymes including enolase, glyceraldehyde-3-phosphate dehydrogenase and phosphoglycerate kinase. Since our analyses detect the protein-nucleic acid conjugate, we can exclude the possibility that these proteins had associated with the RNA following lysis, or were otherwise non-covalently bound. However, the range and number of proteins identified and the prevalence of RNA association with unstructured regions that lack any domain annotation would seem to argue against specific, conserved interactions. It currently appears likely that, despite binding RNA *in vivo*, most of these proteins do not have defined functions in RNA metabolism.

We anticipate that the resources provided here, a protocol to study protein-RNA binding sites at global scale and a list of protein-RNA attachment sites in *S. cerevisiae*, will stimulate systematic and detailed studies of the functionally important protein-RNA interface and may shed light on the hand-over that is envisaged to have occurred during evolution from RNA- to protein-dominated metabolism.

## Experimental Procedures

### Yeast culture

*Saccharomyces cerevisiae* strain BY4741 (*MATa his3*Δ*1 leu2Δ0 met15Δ0 ura3Δ0*) was used for all experiments except SILAC labeling, for which a derivative of BY4742 was used (*MATalpha, his3*Δ*1, leu2Δ0, lys2Δ0, ura3Δ0*) carrying a *LYS2* deletion to block endogenous production of lysine. Non-crosslinked samples were grown is SD medium with 2% glucose and complete supplement mix minus tryptophan and lysine (MP Biomedicals) with unlabeled lysine. Crosslinked samples were grown in the presence of [^13^C_6_] lysine.

### UV crosslinking

Actively growing yeast cultures were crosslinked as described (Granneman et al., 2010). Cells were collected by centrifugation, washed once with PBS and pelleted again. The final pellet was resuspended in an equal volume of ice-cold PBS and pipetted drop-by-drop into liquid nitrogen to snap-freeze cells. Frozen cells were stored at −80°C.

### Cell lysis

Cells were lysed using a 6770 Freezer/Mill Cryogenic Mill (SPEX^®^ SamplePrep) using the manufacturer’s recommendations. For total RNA extraction, powdered yeast was resuspended in 3 volumes of denaturing lysis buffer (7 M urea, 2 M thiourea, 50 mM TRIS/HCl pH 8, 20 mM EDTA, 10 mM DTT), RNA was extracted by vortexing at room temperature after which the lysate was clarified by centrifugation and either processed immediately or stored at −80 °C.

For poly(A)^+^ RNA enrichment, powdered yeast was resuspended in 5 vol lysis buffer (50 mM TRIS/HCl (pH 7.5), 20 mM EDTA, 150 mM NaCl, 1 mM DTT), vortexed five times for 1 min at room temperature with a 1 min incubation at RT after which lysate was clarified by centrifugation and processed immediately.

### Protein digestion

Reduced cysteines were blocked with 20 mM iodoacetamide for 1-2 h at room temperature in darkness. For LysC protein digestion, LysC was added in 1:100 LysC:protein w/w ratio and proteins were digested for 16 h at room temperature. Trypsin digestion was performed under similar conditions, except when urea was used then its concentration was reduced to less than 2 M before digestion.

### RNA purification

Total RNA was purified using CsTFA (GE Healthcare) density centrifugation. RNA was either pelleted through a CsTFA cushion (Zhang, Chen et al., 2003) or separated on a CsTFA step gradient (Smale & Sasse, 1992).

Poly(A)^+^ RNA was enriched using oligo(dT)-cellulose (GE Healthcare) according to the manufacturer’s protocol. RNeasy Mini and Midi columns (QIAGEN) were used according to manufacturer’s protocols.

### RNA hydrolysis

RNA was nuclease-digested as follows: RNA was denatured by heating for 2 min at 100 °C, followed by quick cooling on ice. P1 nuclease (Sigma) was added in 1:500 nuclease RNA w/w ratio and RNA digested for 2 hours at 50 °C. For snake venom phosphodiesterase (Sigma) digestion, TRIS/HCl (pH 8.5) was added to final concentration of 50 mM, phosphodiesterase was added (0.002 U per 1 mg of RNA) and digestion completed for 16 hours at 37 °C.

### Peptide clean-up

Peptides were fractionated on SCX-StageTips as described in (Ishihama, Rappsilber et al., 2006, Rappsilber, Mann et al., 2007).

Mixed mode chromatography (Phillips, Williamson et al., 2010) was performed using Promix MP 250 mm x 2.1 mm id, pore size 300 Å (SIELC, USA) on an Ultimate 3000 HPLC (Dionex,UK) and the following chromatographic buffer conditions: buffer A 10 mM ammonium acetate pH 6.3, 1 M NaCl, buffer B 10 mM ammonium acetate pH 6.3, 80% acetonitrile. Linear gradient elution was performed: 20-80% over 22 min, followed by 100% B over 2 min at a flow rate of 0.2 mL/min. Chromatograms were recorded using UV absorbance at a wavelength of 260 and 280 nm. Fractions were collected at 1 min time interval and desalted using C18-StageTips (Rappsilber, Ishihama et al., 2003, Rappsilber et al., 2007).

### LC-MS/MS analysis

All samples were analyzed on an LTQ Orbitrap Velos or Q Exactive mass spectrometer (Thermo Fisher Scientific), using Ultimate 3000 RSLCnano (Thermo Fisher Scientific) or nanoACQUITY (Waters) LC as the front-end. Peptides were loaded onto 75 µm × 300 mm fused silica emitters (New Objective), packed in-house with Reprosil-Pur C18-AQ 3 µm particles (Dr Maisch, Germany) as described in (Ishihama, Rappsilber et al., 2002) and separated using linear gradient from 2% to 40% of solvent B (solvent A: MilliQ H2O/0.1% formic acid, solvent B: 80% acetonitrile/0.1% formic acid) with a flow-rate of 200 nL/min. Both LTQ Orbitrap Velos and Q Exactive mass spectrometer were operated in data-dependent mode, with dynamic exclusion enabled.

### Data analysis

Peak lists for were generated by MaxQuant 1.4.1.2 (Cox & Mann, 2008).

An open modification search approach using xiOMS (ERI, Edinburgh) was used to identify any unknown modification masses while a targeted modification search using xiTMS (ERI, Edinburgh) was used to analyze data with pre-determined modifications masses (corresponding to RNA crosslink products) but with no amino acid specificity. Parameters used for searches are described in Supplemental Experimental Procedures.

In addition, data were processed with MaxQuant 1.4.1.2 (Cox & Mann, 2008), with following parameters: *S. cerevisiae* UniProtKB sequence database (downloaded on 09.09.2011); protease specificity was Trypsin/P; two missed cleavages were allowed; carbamidomethylation was set as a fixed modification; oxidation (M), N-acetylation (protein) and UMP (YFWILVAK) were set as variable modifications; all FDR levels were set to 0.01.

Protein domain analysis, alignment and visualization was done with pepWizRd, a data analysis application developed in R using the packages shiny, bio3d, CHNOSZ, RSVGTipsDevice and RColorBrewer (Spitzer et al., in preparation).

The mass spectrometry proteomics data have been deposited to the ProteomeXchange Consortium via the PRIDE partner repository with the dataset identifier PXD011511.

## Author Contributions

L.P., D.T. and J.R. designed research; L.P. and S.W. and E.P. performed biochemistry; L.P. and S.W. performed and analyzed MS data, L.P., M.S. and L.F. performed bioinformatics; D.T. and J.R. analyzed data and wrote the paper.

## Acknowledgements

The Wellcome Trust generously funded this work through a Senior Research Fellowship to J.R. (084229), a Principle Research Fellowship to D.T. (077248), a Centre core grant (092076) and an instrument grant (091020), with additional support from the Wellcome Trust/Edinburgh University via the Institutional Strategic Support Fund. L.P. was supported by a Marie Curie Intra European Fellowship within the 7th European Community Framework Programme and by a Estonian Research Council grant (PUT626).

